# Microrheology for Hi-C Data Reveals the Spectrum of the Dynamic 3D Genome Organization

**DOI:** 10.1101/756965

**Authors:** Soya Shinkai, Takeshi Sugawara, Hisashi Miura, Ichiro Hiratani, Shuichi Onami

## Abstract

The 1-dimensional information of genomic DNA is hierarchically packed inside the eukaryotic cell nucleus and organized in 3-dimensional (3D) space. Genome-wide chromosome conformation capture (Hi-C) methods have uncovered the 3D genome organization and revealed multiscale chromatin domains of compartments and topologically associating domains (TADs). Moreover, single-nucleosome live-cell imaging experiments have revealed the dynamic organization of chromatin domains caused by stochastic thermal fluctuations. However, the mechanism underlying the dynamic regulation of such hierarchical and structural chromatin units within the micro-scale thermal medium remains unclear. Microrheology is a way to measure dynamic viscoelastic properties coupling between thermal microenvironment and mechanical response. Here, we propose, to our knowledge, a new microrheology for Hi-C data to analyze the compliance property as a barometer of rigidness and flexibility of genomic regions along with the time evolution. Our method allows conversion of a Hi-C matrix into the spectrum of the rheological property along the genomic coordinate of a single chromosome. To demonstrate the technique, we analyzed Hi-C data during the neural differentiation of mouse embryonic stem cells. We found that TAD boundaries behave as more rigid nodes than the intra-TAD region. The spectrum clearly shows the rheological property of the dynamic chromatin domain formations at an individual time scale. Furthermore, we characterized the appearance of synchronous and liquid-like inter-compartment interactions in differentiated cells. Together, our microrheology provides physical insights revealing the dynamic 3D genome organization from Hi-C data.

**SIGNIFICANCE:** Genomic DNA is hierarchically packed inside the eukaryotic cell nucleus, and the genome organization in 3D contributes to proper genome functions at the multiscale chromatin domains. Although thermal fluctuations inevitably drive movements of the genome molecules in the micro-scale cell environment, there is no method, as yet, to quantify such dynamic 3D genome organization of hierarchical and structural chromatin units. Here, we describe a method to calculate rheological properties as barometers of flexibility and liquid-like behavior of genomic regions. We show that biologically relevant boundaries between chromatin domains are more rigid than the inside at a particular time scale. Our method allows interpretation of static and population-averaged genome conformation data as dynamic and hierarchical 3D genome picture.

## INTRODUCTION

In eukaryotes, the 1-dimensional information of genomic DNA is spatiotemporally organized inside the cell nucleus, which is only a few microns size (1, 2). Dynamic orchestration of genomic regulatory elements in 3-dimensional (3D) space contributes to proper expression of genes. Genome-wide chromosome conformation capture (Hi-C) and related methods have revealed that chromatin hierarchically forms various sized genomic domains such as topologically associating domains (TADs) at the sub-megabase scale and A/B compartments at the megabase scale as a functional and cooperative unit (2–5). These hierarchical folding patterns depend on cell types and states during cell differentiation (6–8). While Hi-C experiments require fixed cells and Hi-C data make sense in population average, the tracking of single nucleosomes by single-molecule and super-resolution live-cell imaging experiments has revealed the dynamic organization of chromatin domains in single cells (9–12). The dynamic property is just like a “polymer melt” state and reveals liquid-like behavior (13, 14). However, the relation of the liquid-like behavior of chromatin to its hierarchical 3D genome organization remains poorly understood.

Thermal fluctuations are dominant and cause Brownian motion in a micro-scale medium as well as within the cell environment. The stochastic dynamics of the Brownian particle can be described by the generalized Langevin equation (GLE), which is formulated from micro-mechanics with the aid of projection methods (15–17). The formalism of microrheology was developed based on GLE (18); the generalized Stokes-Einstein relation (GSER) allows calculation of linear viscoelastic quantities from the mean-squared displacement (MSD) of tracer particles in a complex fluid. Microrheology to measure elastic or viscous properties as mechanical responses in a micro-scale complex fluid has been verified for over two decades (18, 19). Besides, bio-microrheology, the study of deformation and flow of biological materials at small length scales, has revealed the nature of the dynamic coupling between cell microenvironment and mechanical response (20–22).

Recently, the quantitative significance of Hi-C contact matrix data was elucidated mathematically by several independent groups (23–25). We developed a polymer modeling and a simulation method, called PHi-C (Polymer dynamics deciphered from Hi-C data), to decipher Hi-C data into polymer dynamics (25). In the mathematical formalism, we found a one-to-one correspondence between a Hi-C contact matrix and an interaction matrix of the polymer model. Once an optimal interaction matrix of the polymer model to an input Hi-C matrix data is obtained, the method allows calculation of not only dynamic information such as the MSD of a modeled genomic region, but also conformations of the polymer model. Therefore, integrating PHi-C with GSER can provide a new microrheology for Hi-C data to characterize the viscoelastic properties of the modeled chromosome dynamics.

Here, we demonstrate that our microrheology converts a Hi-C matrix into the spectrum of the complex compliance *J*^*^ (*ω*) as a barometer of the flexibility of a modeled genomic region. As *ω* represents a frequency of oscillatory shear stress in the rheology formalism (26), the inverse *t* = *ω*^−1^ corresponds to a time and *J*^*^ (*ω*) should include dynamic information along the time. Thus, the converted spectrum of the complex compliance describes how an individual genomic region as a part of the modeled polymer flexibly deforms at a specific time scale. We show that rigid boundaries in a profile of the complex compliance along a genomic coordinate at a time scale statistically characterize TAD boundaries. Moreover, by applying Hi-C data during cell differentiation, we find the appearance of synchronous and liquid-like inter-compartment interactions in differentiated cells. Taken together, our microrheology enables us to interpret Hi-C data as a physical picture regarding the dynamic 3D genome organization.

## MATERIALS AND METHODS

### Theory of microrheology to convert Hi-C data into complex compliance

For Hi-C matrix data with an appropriate binned resolution, a single chromosome was modeled by the polymer network model (25), which consists of *N* monomers. The attractive or repulsive interaction between two monomers was assumed as a linear force proportional to the displacement between the two monomers. Therefore, the physical interactions of all pairs can be described as an *N* × *N* matrix K = *K*_*ij*_. The positive values of K stand for elastic forces between two monomers, and the model formally resembles the Gaussian network model (27–29). Here, the negative values are acceptable as repulsive forces in the polymer network model. This assumption is a unique point of our modeling and different from recently developed similar polymer modeling (23, 24). As long as the Laplacian matrix of K is positive-semidefinite, the polymer organization does not break down (25).

PHi-C (https://github.com/soyashinkai/PHi-C) is a simulation tool to decipher Hi-C data based on the mathematical formalism of the polymer network model, in which a normalized contact matrix C = (*C*_*ij*_) is connected on one-to-one correspondence with the normalized interaction matrix 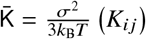 (25). Here σ represents the contact distance, *k*_B_ is the Boltzmann constant, and *T* is the temperature of the environment. The PHi-C optimization procedure allows for extracting an optimal normalized interaction matrix 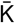 from an input normalized contact matrix C (Fig. 1).

**Figure 1:**
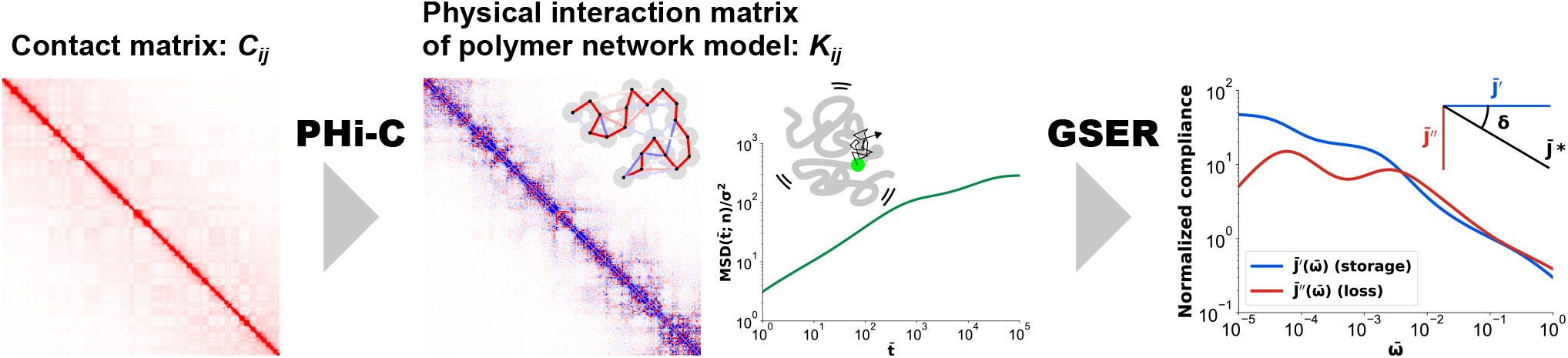
Microrheology pipeline for Hi-C data using PHi-C and GSER. A normalized contact matrix *C*_*ij*_ as input data is deciphered into a normalized physical interaction matrix of the polymer network model 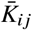 through PHi-C optimization. Red and blue colors in the bins of 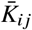 represent the intensity of the attractive and repulsive interactions, respectively. The MSD of the *n*-th monomer in the polymer model is analytically calculated from the interaction matrix 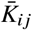. Finally, the GSER allows to obtain the normalized complex compliance 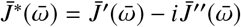. The ratio 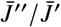 corresponds to the loss tangent, tan *δ*, where *δ* is the phase angle.

In theory, the physical interaction matrix K includes all information with respect to not only dynamics but also conformations in thermal equilibrium. The matrix K can be converted into the Laplacian matrix L = D − K to characterize the properties of the network, where the degree matrix is defined by D = diag (*D*_0_, *D*_1_, …, *D*_*N*−1_) and 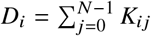. As the Laplacian matrix L is real symmetric, L is diagonalizable. Furthermore, the *N* eigenvalues satisfy 0 = *λ*_0_ < *λ*_1_ ≤ *λ*_2_ ≤ … ≤ *λ*_*N*−1_ as long as L is positive-semidefinite, and there is an orthogonal matrix Q such that Q^T^LQ = diag (*λ*_0_, *λ*_1_, …, *λ*_*N*−1_). Then, the MSD of the *n*-th monomer within the modeled single chromosome can be written as

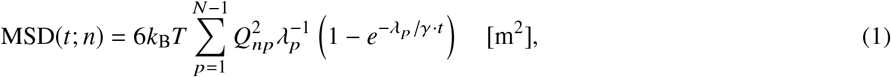

where *γ* is the friction coefficient of the monomers, and *t* represents actual time.

The GSER connects the MSD to the complex shear modulus in the Laplace domain (18), with the inertia being neglected, 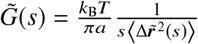 [Pa], and the complex compliance is defined as the inverse of the complex shear modulus (26),

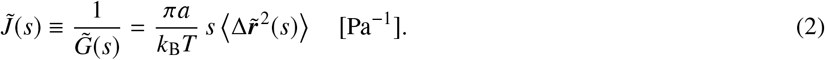

Here, we employed the MSD of a modeled genomic monomer derived by PHi-C in order to consider the dynamic viscoelastic properties of the genome itself within a single chromosome. Calculating the Laplace transformation of Eq. 1, the complex compliance of the *n*-th monomer within a single chromosome was expressed by using the eigenvalues 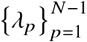 and the orthogonal matrix Q as

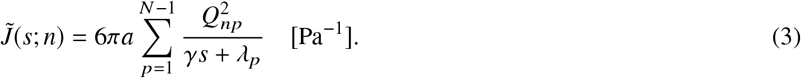

The matrix one-to-one correspondence between C and 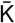 in PHi-C is formally closed as dimensionless quantities. Therefore, the above complex compliance with the physical unit Pa^−1^ should be normalized. The eigenvalues of the normalized Laplacian matrix 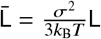 were normalized as 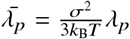. Moreover, using the normalized inverse time 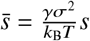, we could rewrite Eq. 3 as

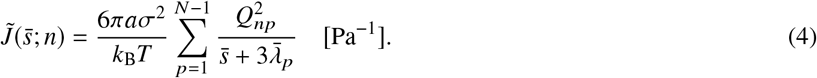

In general, the complex compliance was defined by the Fourier-Laplace transformation, replacing 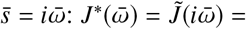 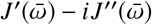, where 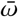, *J*′ and *J*′′ are the normalized frequency, the storage and loss compliances, respectively. Thus, the normalized complex compliance was derived from the result of PHi-C with no reference to physical parameters,

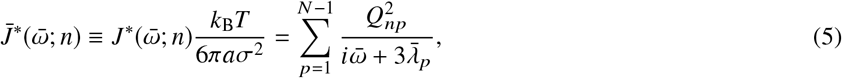

and the normalized storage and loss compliances can be written as

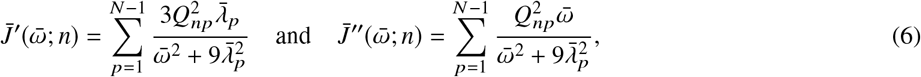

respectively. The ratio of 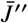 to 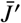 is called the loss tangent,

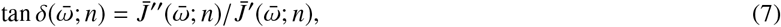

where *δ* represents the phase angle between stress and strain (26).

### Hi-C data of mouse embryonic stem cells

We used Hi-C data of mouse embryonic stem (ES) cells during neural differentiation by Bonev et al. (8) (GEO: GSE96107) on the Hi-C data archive of Juicebox (30). Using Juicer Tools (30), we extracted the Hi-C matrix data with KR normalization and at 250-kb resolution from *.hic* files, and calculated the eigenvector. The extracted Hi-C matrix data were analyzed through the PHi-C pipeline (25).

### Analysis of the distance between TAD boundaries and the genomic positions of the troughs

The analysis was performed as previously described with small modifications (31, 32). We used TAD boundary data on chromosome 6 of ES cells, defined by Bonev et al. (2). To generate the control trough list, the genomic positions of the troughs at each frequency (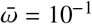, 10^−2^ and 10^−3^) were randomly permutated at 250-kb genomic bin resolution on chromosome 6 excluding the centromeric region (0–3 Mb). Then, cumulative probabilities of the overlap between the lists of TAD boundaries and the troughs (original or randomly permutated) at a given frequency were computed based on their nearest distance.

## RESULTS

### Spectrum of complex compliance during mouse neural differentiation

To calculate rheological properties of the 3D genome organization during neural differentiation of mouse ES cells by using PHi-C and GSER (Fig. 1), we analyzed deep coverage Hi-C data of mouse ES cells, neural progenitors (NPCs), and cortical neurons (CNs) (8). First, we performed the PHi-C optimization procedure for the 3–149.75-Mb genomic region of chromosome 6 to obtain an optimal normalized interaction matrix 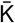 of the polymer network model. The optimized contact matrices for ES, NPC, and CN cells showed very high Pearson’s correlations (0.979, 0.978, and 0.959, respectively) between the Hi-C matrix and the optimized contact matrix (Fig. 2).

**Figure 2:**
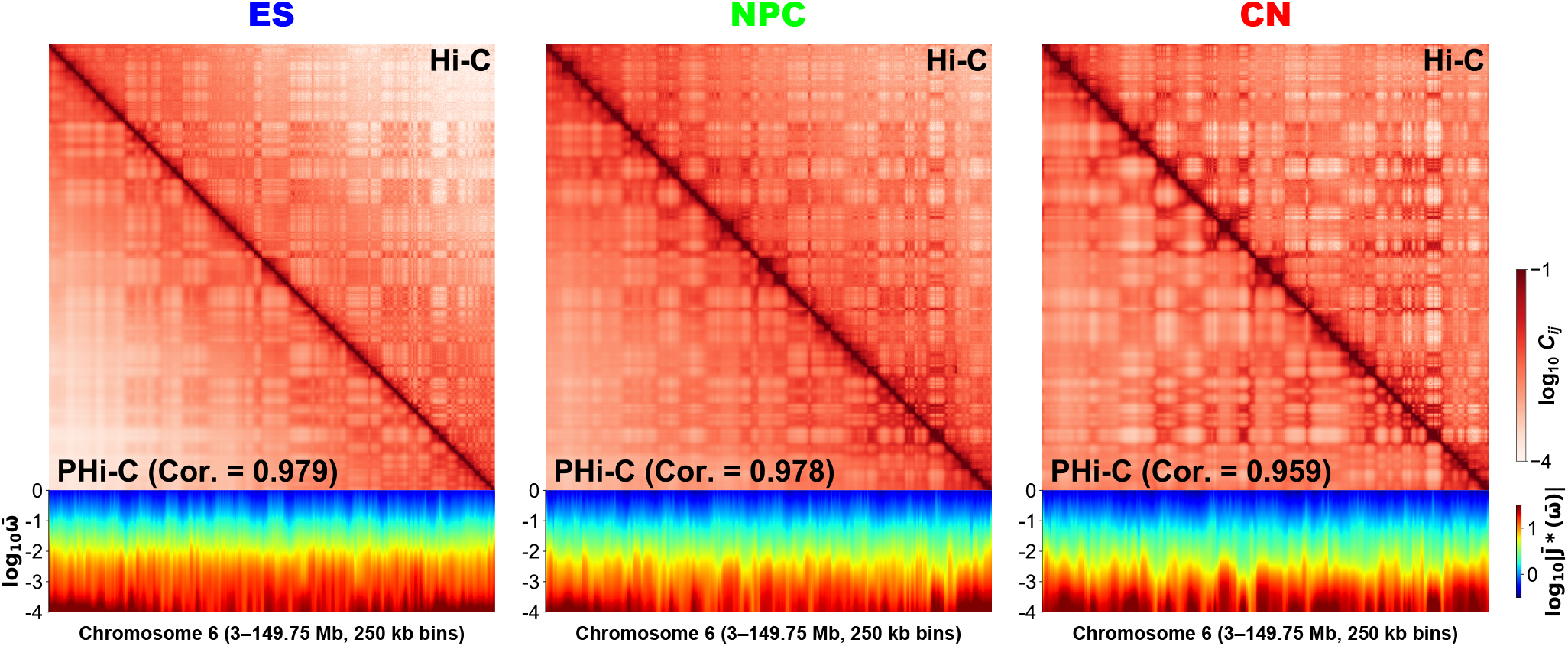
Contact matrices and spectra of the normalized complex compliance 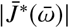 for chromosome 6 during mouse neural differentiation of embryonic stem (ES) cell (Left), neural progenitor (NPC) (Middle), and cortical neuron (CN) (Right) at 250-kb resolution. Upper right and lower left elements in the matrix correspond to the normalized Hi-C contact probabilities and the optimized ones by PHi-C, respectively. Along the genomic coordinate and the logarithmic frequency 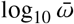, a spectrum of 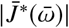 is depicted as a heat-map.

Using Eqs. 5 and 6, we calculated the normalized complex compliance 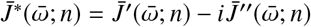 as a barometer of flexibility for an oscillatory stress with the normalized frequency 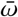. In addition, the parameter *n*(= 0, 1, …, 586) represents a genomic region (3 + 0.25 × *n*)−(3 + 0.25 × (*n* + 1))-Mb. Thus, we obtained a spectrum of 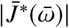 along the genomic coordinate (Fig. 2). According to changes in the Hi-C patterns during mouse neural differentiation, the spectra of 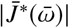 also showed the different patterns.

In general, at a fixed frequency *ω* [Hz], rheological quantities express the viscoelastic response to the periodic perturbation with *ω*. In other words, these quantities include the dynamic information on the viscoelastic mobility at the time scale *t* = *ω*^−1^ [s]. Therefore, the spectrum in Fig. 2 should show dynamic hierarchy in a single chromosome as we can infer a hierarchical 3D genome organization from a Hi-C contact pattern.

### TAD boundaries are more rigid as nodes than intra-TAD sequences

To understand what the values of the complex compliance 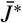 reveal along the genomic coordinate, we plotted 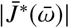 at 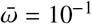, 10^−2^ and 10^−3^ focusing on the 50–100 Mb region of the chromosome 6 of ES cells (Fig. 3A). As we can see variously sized triangles corresponding to chromatin domains such as TADs and compartments on the Hi-C contact pattern, the shapes of 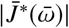 displayed peaks and troughs. We found that the troughs are located near the TAD boundaries at each frequency 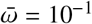, 10^−2^ and 10^−3^ with a statistical test (Fig. 3A and Fig. S1), suggesting that TAD boundaries are characterized as more rigid nodes than the intra-TAD sequence region according to the time scale 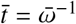. Therefore, the vertical stripe pattern on the spectrum in Fig. 2 shows the variation of the peaks and troughs depending on the frequency.

**Figure 3:**
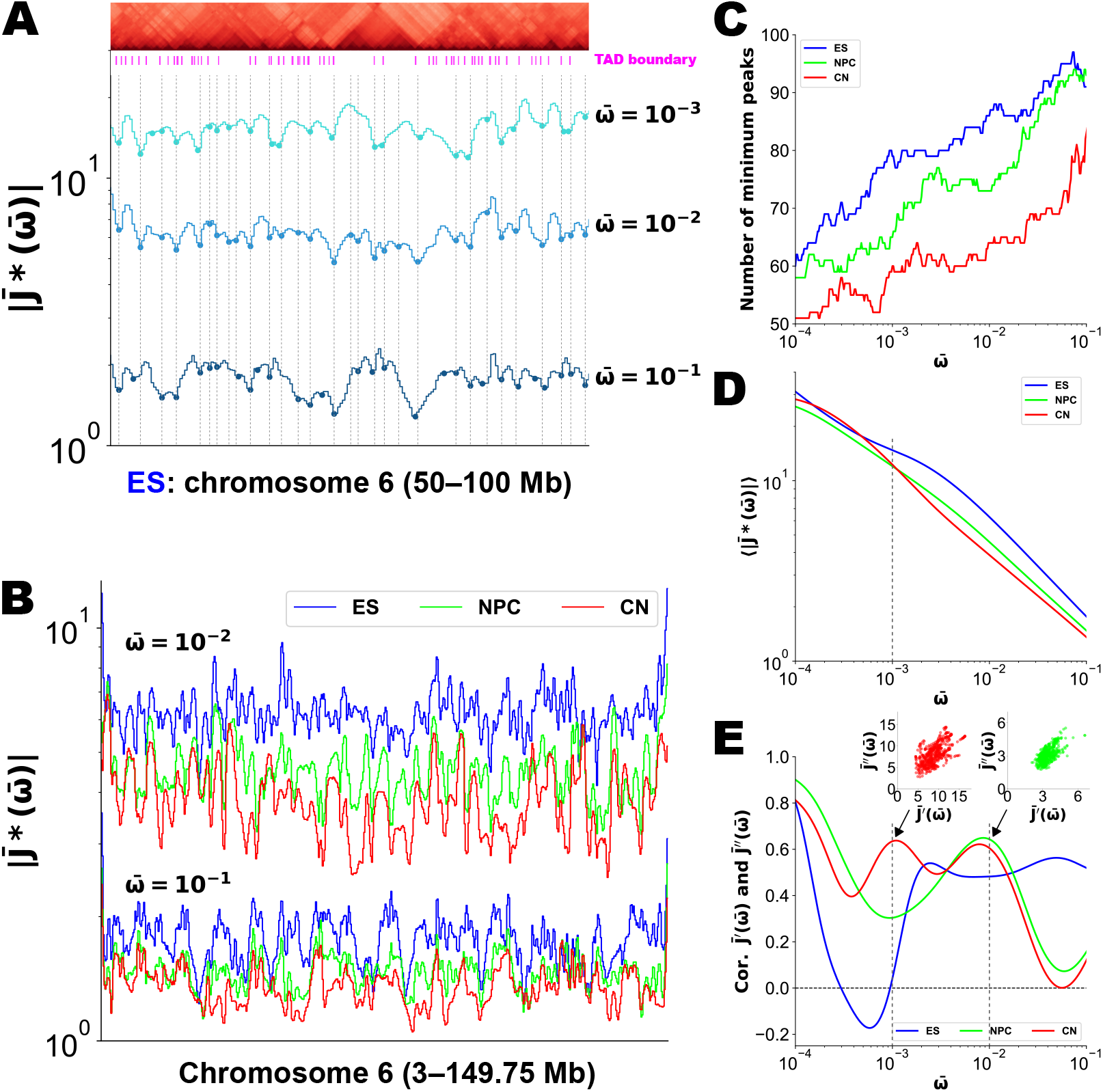
Complex compliance characterizes chromatin domain boundaries and insides during mouse neural differentiation. (A) 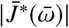 at 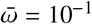, 10^−2^ and 10^−3^, and a cropped Hi-C contact matrix for the 50–100 Mb region of embryonic stem (ES) chromosome 6 with a TAD boundary profile (8). The vertical dashed lines correspond to the genomic position at the troughs of 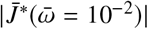. (B) 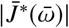 at 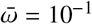 and 10^−2^ for ES, neural progenitor (NPC), and cortical neuron (CN) chromosome 6. (C) The number of minimum peaks as the troughs on 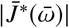 for ES, NPC, and CN chromosome 6. (D) Average values of 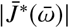 along chromosome 6 for ES, NPC, and CN. (E) Pearson’s correlation for the storage and loss compliances, 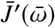 and 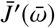 within ES, NPC, and CN chromosome 6. Scatter plots at 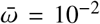 and 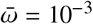 for NPC and CN are displayed, respectively.

### Dynamic and hierarchical changes of chromosome rigidness during cell differentiation

The peaks and troughs in the shape of 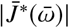 were also observed for chromosome 6 of ES cells, NPCs, and CNs (Fig. 3B). The number of troughs decreased as time 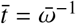 increased (Fig. 3C). This indicated that dynamic and hierarchical compartmentalization with domain fusions occur in the 3D genome organization according to the time evolution. For example, intra-TAD dynamics is dominant until 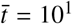, then inter-TAD communications and fusions of TADs arise up to 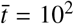 in intra-compartments, and inter-compartment interactions occur over 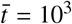. Besides, the numbers of troughs for ES, NPC, and CN were ordered, revealing that the number of chromatin domains as chromosome structural units decreases depending on an individual frequency, and the rearrangement occurs during cell development. Furthermore, the average values of 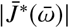 were also ordered until the time 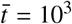 (Fig. 3D), suggesting that the physical property of chromosomes becomes rigid and less flexible during cell differentiation.

The complex compliance 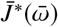 is divided into the storage and loss components (Eq. 6), which are not independent of each other in rheological relations (26). How these compliances correlate within ES, NPC, and CN chromosome 6 at a fixed frequency 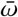 is shown in Fig. 3E. For ES cells, broad positive correlations were detected in the frequency region 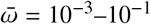. A high positive correlation indicates that the elastic and viscous responses described by 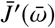 and 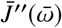 synchronize within a chromosome to the periodic stress with a frequency 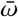. Therefore, a positive peak of the correlation indicates a characteristic time 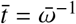 of the synchronous response. For NPCs and CNs, a broad region showing high correlations was not observed. Instead, some peaks were observed around 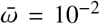 for NPC and 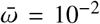 and 10^−3^ for CN. Note here that the peaks around 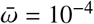 are excepted because the response to the extremely slow fluctuation corresponds to a whole movement of the chromosome. As we could confirm that the checkerboard pattern corresponding to the A/B compartment organization in Fig. 2 gradually becomes dense during cell differentiation, the correlations for ES, NPC and CN at 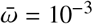 are ordered, suggesting that the time scale 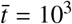 must be related to inter-compartment organization.

### Synchronous and liquid-like inter-compartment interactions appear in differentiated cells

Finally, we asked whether the microrheological property revealed by our method relates to the A/B compartment organization. We depicted spectra of the loss tangent 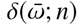 (see Eq. 7) with the eigenvector to characterize the compartment organization (3) during differentiation in Fig. 4A. In general, positive and negative values of the logarithmic loss tangent indicate solid-like elasticity and liquid-like viscosity, with lower and higher energy dissipation, respectively.

**Figure 4:**
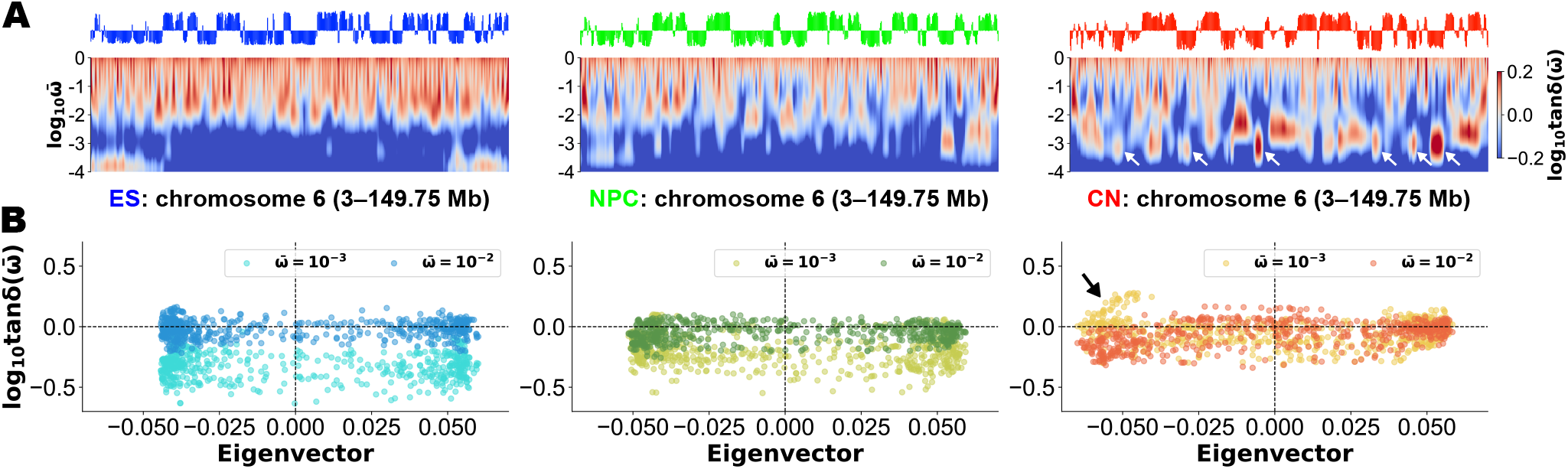
(A) Eigenvectors and spectra of the loss tangent 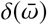 for embryonic stem (ES) (Left), neural progenitor (NPC) (Middle), and cortical neuron (CN) (Right) chromosome 6 at 250-kb resolution. Along the genomic coordinate and the logarithmic frequency 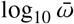, a spectrum of 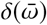 is depicted as a heat-map. White arrows for CN indicate definite “island” regions around 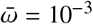 with negative eigenvectors. (B) Scatter plots between the logarithmic loss tangent and the eigenvalue for 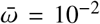 and 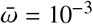. For CN, the black arrow indicates the appearance of the “islands” in (A).

For ES cells, the spectrum showed that viscous behavior with higher energy dissipation is dominant in the frequency 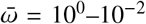 and the shape at a fixed frequency has peaks and troughs. However, the positive and negative values of the logarithmic loss tangent for 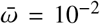 did not correlate with the eigenvalues (Fig. 4B; Left), implying that the loss tangent cannot distinguish the active compartments from the inactive compartments. In addition, the scatter plot for 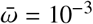 dropped horizontally, compared with that for 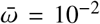. For NPCs, we could confirm a different pattern in the spectrum, although the scatter plots for 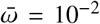 and 10^−3^ showed an overall decrease similar to that for ES cells (Fig. 4A and 4B; Middle).

In contrast, the spectrum for CNs revealed a characteristic pattern of isolated regions like “islands” with the positive values (Fig. 4A; Right). Interestingly, some islands synchronously appeared around the frequency 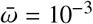 belonging to the genomic region with negative eigenvalues. The synchronous appearance was also confirmed in the scatter plots for 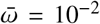 and 10^−3^ (Fig. 4B; Right), where the solid-like elastic property (tan *δ* < 1) for 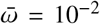 turns into the liquid-like viscous property (tan *δ* > 1) for 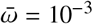.

## DISCUSSION

In this study, we revealed the rheological information of the dynamic 3D genome organization as a spectrum by integrating the theoretical MSD by PHi-C and GSER. On the spectrum, the differences in the compliance along a genomic coordinate at a particular time scale characterized the boundaries and insides of chromatin domains: especially, TAD boundaries were found to be more rigid than the intra-TAD region. During cell differentiation, we quantitatively estimated increasing rigidity of the chromosome, with dynamic organization of chromatin domains with time evolution.

Particularly, our microrheology method allows for interpreting static and population-averaged Hi-C matrix data as a dynamic and hierarchical 3D genome picture. Definite boundaries with higher rigidness at a particular time scale 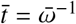 characterize an individual triangle region corresponding to a chromatin domain such as a TAD or a compartment on a Hi-C pattern. In other words, the appearance of a chromatin domain as a functional unit of the 3D genome organization correlates to a specific time scale with the dynamics. In addition, organization of an inter-domain interaction is relatively slow, and synchronous and liquid-like inter-compartment interactions were notable in the mouse neural differentiated cell. Therefore, the spectrum revealed by our microrheology method explicitly shows the hierarchical 4D genome organization beyond schematic and static 3D genome pictures on paper.

Large-scale numerical simulations for modeling the interphase cell nucleus environment have revealed viscoelastic properties based on GSER (33). As microrheology experiments have tracked the motion of tracer particles, the authors simulated the viscoelastic response of dispersed Brownian particles in polymer solutions and found a caged effect at short times large particles. In the current study, our microrheology method did not focus on the tracer particles, but rather on modeled chromosome dynamics itself by calculating the theoretical MSD and using GSER. Thus, we could characterize the rheological properties of a chromosome along the genome coordinate. Moreover, we used the complex compliance *J*^*^(*ω*) as a barometer of flexibility, but we can convert it into the complex modulus *G*^*^(*ω*) and the dynamic viscosity *η*^*^(*ω*) by the relationship *G*^*^(*ω*) = *i*(*ω*)*η*^*^(*ω*) = 1/*J*^*^(*ω*) (26).

Although the PHi-C optimization procedure can return a high correlation value between an input Hi-C and the optimized contact matrices, the correlation value depends on not only the quality and matrix normalization of the Hi-C data, but also the computational time of the PHi-C optimization. Besides, although the theoretical MSD result was not comprehensively verified by experiments, the MSD curves theoretically derived by PHi-C for *Nanog* and *Oct4* loci of ES cells were partially consistent with the genomic movements in a live-cell imaging experiment (25, 34). Under a high correlation value by the PHi-C optimization and assuming that a chromosome modeled by the polymer network model is in thermal equilibrium, our microrheology method is reliable and competent.

In this study, we focused on the rheological properties of a specific chromosome during cell development, but we could not elucidate a relationship between the physical properties of chromatin and gene expression, i.e. how physical rigidness or flexibility of a specific genomic region including some loci might relate to the gene expression levels. Our findings on TAD boundaries as more rigid nodes would suggest the molecular mechanism regulating the physical rigidness of the boundaries.

## CONCLUSION

In summary, the microrheological spectrum generated from Hi-C data describes the dynamic change of rigidness and flexibility in an individual genomic region as well as during cell differentiation. The appearance of chromatin domain formations and inter-domain interactions requires an individual time scale. Our microrheology method opens the possibility of interpreting Hi-C data as information of the dynamically hierarchical genome organization.

## AUTHOR CONTRIBUTIONS

S.S., T.S., and S.O. designed the research. S.O. supervised the study. S.S. carried out analytical calculations and PHi-C analysis, and analyzed the data. H.M. and I.H. performed TAD boundary analysis. S.S. wrote the article.

## ACKNOWLEDGMENTS

This work was supported by JSPS KAKENHI Grant Numbers JP18H04720 to S.S., JP17K15050 to T.S., JP18H05530 to I.H., and JP18H05412 to S.O.; Core Research for Evolutionary Science and Technology (CREST) Grant Number JPMJCR1511, Japan Science and Technology Agency (JST) to S.O.

