## Supplementary Figure S1 for "Microrheology for Hi-C Data Reveals the Spectrum of the Dynamic 3D Genome Organization"

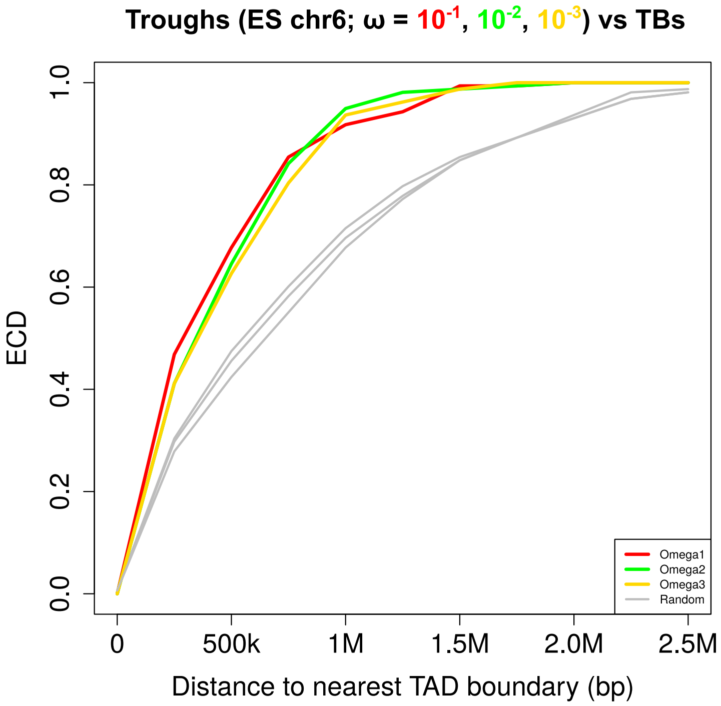


**Supplementary Figure S1: Cumulative probability analysis for TAD boundaries and the genomic positions of the troughs.** A cumulative probability analysis of the distance between the trough positions and the nearest TAD boundaries on chromosome 6 in mouse embryonic stem (ES) cells. Red, green, gold, and grey represent the troughs from the frequency $\bar{\omega} = 10^{-1}$, $10^{-2}$ and $10^{-3}$, and randomly permutated (control), respectively, with the former three significantly closer to the TAD boundaries than control ($p = 9.42 \times 10^{-6}$, $9.44 \times 10^{-6}$, $1.04 \times 10^{-5}$, respectively, by a two-sided Wilcoxon rank sum test).
